# Genome-wide Association Study for Number of Vertebrae in an F2 Large White × Minzhu Population of Pigs

**DOI:** 10.1101/016956

**Authors:** Longchao Zhang, Xin Liu, Jing Liang, Kebin Zhao, Hua Yan, Na Li, Lei Pu, Yuebo Zhang, Huibi Shi, Ligang Wang, Lixian Wang

**Affiliations:** Key Laboratory of Farm Animal Genetic Resources and Germplasm Innovation of Ministry of Agriculture of China, Institute of Animal Science, Chinese Academy of Agricultural Sciences, 100193 Beijing, China

**Keywords:** genome-wide association study, number of vertebrae, pig, SSC7, *TGFB3*

## Abstract

Porcine carcass that is approximately 800 mm long may be expected to have one additional vertebra. Therefore, the number of vertebrae in pigs is an economically important trait. To examine the genetic basis of this trait, we genotyped 593 F2 Large White × Minzhu pigs using the Porcine SNP60K BeadChip. A genome-wide association study identified 39 significant single-nucleotide polymorphisms (SNPs) on the chromosomes SSC1 and SSC7. An 8.82-Mb region that contained all 21 significant SNPs on SSC1 harbored the gene *NR6A1*, previously reported to influence the number of vertebrae in pigs. The remaining 18 significant SNPs on SSC7 were concentrated in a 4.56-Mb region, which was within the quantitative trait loci interval for number of vertebrae. A haplotype sharing analysis refined the locus to a segment of ∼637 Kb. The most significant SNP, SIRI0001067, was contained in this refined region on SSC7 and located in one of the introns of *TGFB3*. As *TGFB3* influences the development of vertebrae in mammalian embryos, the gene may be another strong candidate for the number of vertebrae in pigs.

---

Vertebrae consist of the morphologically differentiated cervical, thoracic, lumbar, sacral, and caudal vertebrae, and the number of these varies in pigs (King and Roberts 1960). Wild boar, the ancestor of domestic pigs, have a uniform 26 cervical–lumbar vertebrae, whereas most Chinese indigenous pig breeds have 26 or 27 vertebrae (Zhang *et al.* 1986). By comparison, Western commercial pig breeds, such as Landrace and Large White, have 27–29 vertebrae (Mikawa *et al.* 2005). The number of cervical–lumbar vertebrae is usually associated with carcass length and is an economically important trait; a greater number of vertebrae in pigs is associated with higher economic value. According to a study by King and Roberts, a carcass that is approximately 800 mm long may be expected to have one additional vertebra (King and Roberts 1960). Understanding the genetic basis for number of vertebrae can offer insights into the mechanism for vertebral developmental in mammals and provide genetic markers to aid molecular breeding of pigs.

Locating quantitative trait loci (QTLs) and gene-mining for number of vertebrae have recently become research interests. Wada *et al.* (2000) fist reported two QTLs on wild boar (*Sus scrofa*) chromosomes SSC1 and SSC2 that are associated with the number of vertebrae. Subsequently, using several different pig populations, two QTLs for the number of vertebrae were mapped to SSC1 and SSC7 (Mikawa *et al.* 2005). Using three F2 experimental families, a fine mapping of the QTL on SSC1 was performed and showed that nuclear receptor subfamily 6, group A, member 1 (*NR6A1*), was a strong candidate gene that appeared to influence the number of vertebrae in pig (Mikawa *et al.* 2007). Gene-mining to the other QTL on SSC7 suggested that the vertebrae development homologue (VRTN) could be a good candidate (Mikawa *et al.* 2011). A genome-wide association study (GWAS), based on the high density of genome-wide single-nucleotide polymorphisms (SNPs), has been a more efficient method to not only identify QTLs but also mine major new genes. Additional methods and different populations are required to identify loci that are associated with the number of ribs and detect good candidate genes. The objectives of this study were to identify SNPs associated with the number of vertebrae using GWAS in an F2 Large White × Minzhu population of pigs.

## MATERIALS AND METHODS

### Ethics statement

All animal procedures were performed according to the guidelines developed by the China Council on Animal Care, and all protocols were approved by the Animal Care and Use Committee at Institute of Animal Science, Chinese Academy of Agricultural Sciences. All efforts were made to minimize the suffering of animals.

### Population and phenotypic data

The F0 population was generated using four Large White boars and 16 Minzhu sows. In the F1 generation, nine boars and 46 sows were selected to mate to produce 598 F2 individuals. All animals in the F2 generation were born in three parties and 94 litters. Each F2 male was castrated 3 days after birth. All animals were maintained under uniform housing conditions and were fed the same fodder. When F2 animals arrived at 240 ± 7 days, they were slaughtered in 52 batches. The numbers of thoracic and lumbar vertebrae were counted from the carcasses.

### Genotyping and quality control

Genomic DNA was extracted using standard methods (Miller *et al.* 1988). The Illumina SNP60K chip for pigs was employed to genotype all individuals. As our previous study (Zhang *et al.* 2014), quality control parameters for single nucleotide polymorphisms included the following: call rate > 90%, minor allele frequency (MAF) > 3%, and Hardy–Weinberg equilibrium (HWE) with a *P*-value > 10^−6^.

### GWAS

The genome-wide rapid association using the mixed model and regression-genomic control approach (Aulchenko *et al.* 2007; Amin *et al.* 2007) was used in the present study. Three fixed effects (i.e., sex, parity, and batch), a random effect (litter), and a covariate (body weight) were used as inputs. DMU (Madsen *et al.* 2006) and GenABEL software (Amin *et al.* 2007) in the R environment were employed to estimate the residuals of traits for each individual and perform the GWAS. After Bonferroni correction, genome-wide significance was accepted at a *p*-value of 0.05/number of SNPs passing quality control (Yang *et al.* 2005).

### Haplotype sharing and linkage disequilibrium analysis

The genotypes of F1 sires were confirmed using marker-assisted segregation analysis (MASS) (Nezer *et al.* 2003). According to the SNP, H3GA0004881, the genotype of each boar was checked from a z-score as the log_10_ likelihood ratio (L_H1_/L_H0_), where L_H1_ supposes that the sire is heterozygous (Qq), and L_H0_ supposes that the sire is homozygous (QQ or qq). Sires were regarded as Qq, if z >2; QQ or qq if z < –2; and of uncertain genotype when 2 > z > –2. The Q-bearing chromosomes of F1 boars and top 10% of F2 population were segregated by MASS. Haplotype sharing analysis was done according to all of significant SNPs on SSC7.

## RESULTS AND DISCUSSION

### Phenotype descriptions

The number of vertebrae in the Large White × Minzhu intercross population may be found in Table 1. The pigs had 24 (n = 1), 25 (n = 9), 26 (n = 244), 27 (n = 296), 28 (n = 42), or 29 (n = 1) vertebrae and the mean number of ribs was 26.63.

**Table 1.** Summary of number of vetebrae in Large White × Minzhu intercross population.

| Total Number of Vertebrae |  |  |  |  |  | Total | Average |
| --- | --- | --- | --- | --- | --- | --- | --- |
| 24 | 25 | 26 | 27 | 28 | 29 |  |  |
| Number of Individuals |  |  |  |  |  |  |  |
| 1 | 9 | 244 | 296 | 42 | 1 | 593 | 26.63 |

### SNPs on SSC1 containing NR6A1

The final data set that passed quality control screening and was used in the analysis contained 47,615 SNPs and came from 585 F2 individuals. The distribution of SNPs after quality control and the average distance between adjacent SNPs on each chromosome are shown in Table 2. The results of the GWAS for number of vertebrae are shown in Table 3. The Manhattan plot obtained from GWAS is shown in Figure 1. On SSC1, there were 21 genome-wide significant SNPs associated with number of vertebrae, within an 8.82-Mb (293.93–302.75 Mb) region. Several previous studies have mapped the QTL for number of vertebrae to a similar interval on SSC1 (Mikawa *et al.* 2005, 2007, 2011; Wada *et al.* 2000; Sato *et al.* 2003; Ren *et al.* 2012). Fine mapping to the QTL showed that *NR6A1* is a strong candidate for the QTL and that the most likely causative mutation is a base substitution, *NR6A1* c.748 C > T, which results in a proline to leucine substitution at codon 192 (Mikawa *et al.* 2007). The strongest signature of selection was observed for a locus on chromosome 1, which includes *NR6A1* (Rubin *et al.* 2012). The distribution of *NR6A1* c.748 C > T in different pig breeds showed that the C and T alleles were almost fixed in wild boars and western commercial breeds, respectively (Burgos *et al.* 2014; Fontanesi *et al.* 2014). All three genotypes are represented in European and Chinese indigenous pig breeds (Burgos *et al.* 2014; Yang *et al.* 2009). In the present study, the most significant SNP, H3GA0004881, was near *NR6A1*, indicating that this gene is a strong candidate for the number of vertebrae in pigs.

**Table 2.** Distribution of SNPs after quality control and the average distances between SNPs on each chromosome^1^.

| Chromosome | No. SNPs | Average distance(kb) |
| --- | --- | --- |
| 1 | 5323 | 59.16 |
| 2 | 2759 | 58.82 |
| 3 | 2228 | 64.69 |
| 4 | 2937 | 48.81 |
| 5 | 1971 | 56.43 |
| 6 | 2620 | 60.02 |
| 7 | 2779 | 48.45 |
| 8 | 2308 | 63.93 |
| 9 | 2668 | 57.52 |
| 10 | 1496 | 52.62 |
| 11 | 1574 | 55.67 |
| 12 | 1308 | 48.49 |
| 13 | 3384 | 64.57 |
| 14 | 3136 | 48.99 |
| 15 | 2361 | 66.68 |
| 16 | 1565 | 55.50 |
| 17 | 1381 | 50.09 |
| 18 | 1133 | 53.74 |
| X | 733 | 195.70 |
| 0<sup>2</sup> | 3951 |  |
| Total | 47615 |  |

<sup>1</sup>Data from *Sus scrofa* Build 10.2
<sup>2</sup>These SNPs are not assigned to any chromosomes.

**Table 3.**
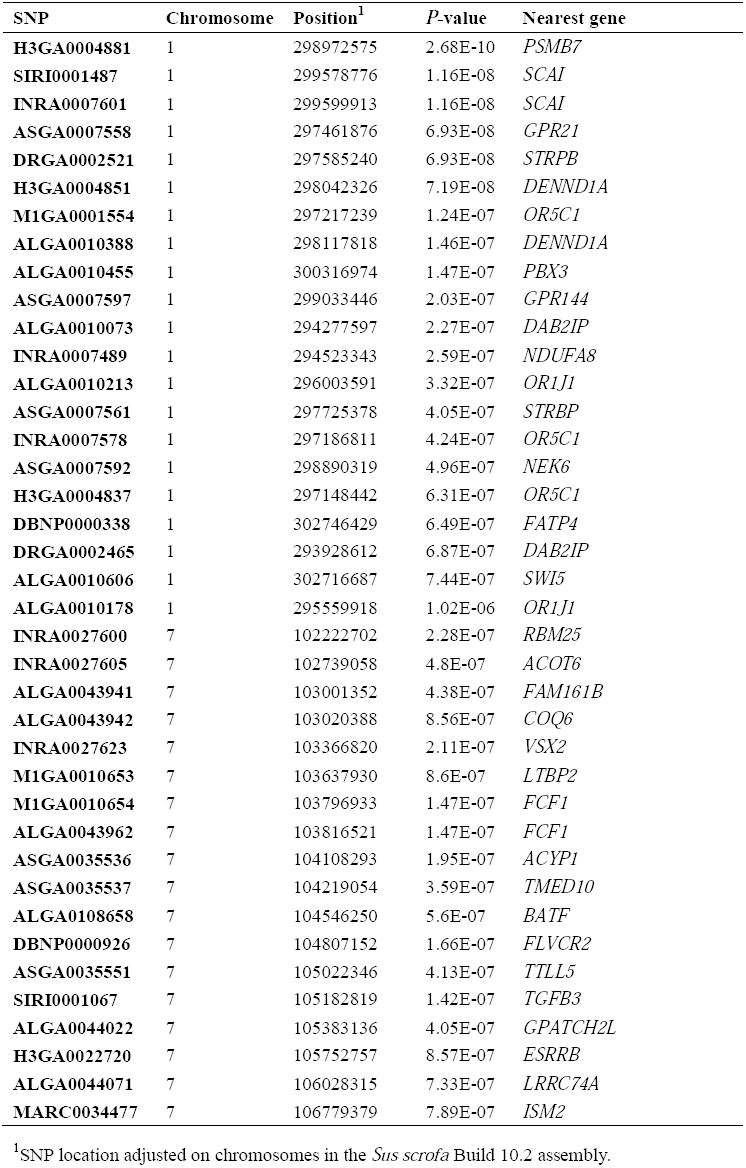
Genome-wide significant SNPs associated with number of vertebrae.

**Figure 1.**
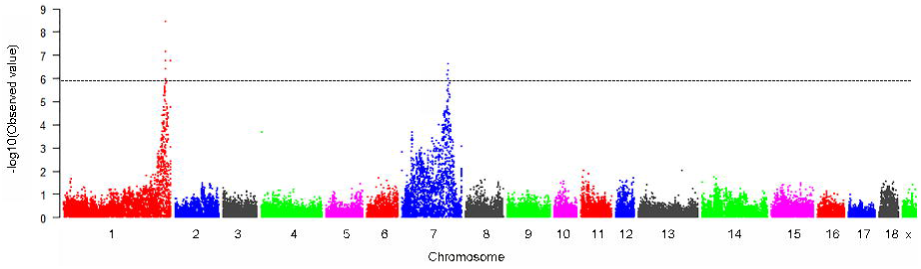
Manhattan plots of genome-wide association study with porcine number of vertebrae. Chromosomes 1-18, and X are shown in different colors. The horizontal line indicates the genome-wide significance level (-log_10_ (1.05E-06)).

### Genome-wide association of SNPs on SSC7 with number of vertebrae

The genome-wide significant SNPs on SSC7 are shown in Table 2. A total of 18 SNPs were significantly associated within a 4.56-Mb region (102.22–106.78 Mb) on SSC7. The most significant SNP was H3GA0004881. Several previous studies have reported similar findings when they mapped the major QTL associated with the number of vertebrae on SSC7 using different populations. This QTL was first reported to be on SSC7 in an F2 Meishan × Duroc resource population (Sato *et al.* 2003). Subsequently, research in several populations derived from crosses between Western pig breeds (Large White, Duroc, Berkshire, and Landrace) and Chinese indigenous pig breeds (Meishan and Jinhua), revealed that the identical QTL was located on SSC7 (Mikawa *et al.* 2005). Moreover, this QTL on SSC7 was repeatedly identified in an F2 population crossed from the commercial breeds Duroc and Pietrain (Edwards *et al.* 2008). Gene-mining in the QTL region revealed that *VRTN* could be a strong candidate gene and the Q/Q homozygotes could increase ∼1 vertebra over the wt/wt (Mikawa *et al.* 2011; Fan *et al.* 2013). In addition, the *VRTN* Q/Q genotype was reported to significantly increase body length by ∼1 cm (Hirose *et al.* 2013). *VRTN* was also located in this GWAS region and could be a good candidate for number of vertebrae in pigs.

### Haplotype sharing analysis to refine the QTL on SSC7

Using the most significant SNP, SIRI0001067, MASS identified three out of nine F1 boars as heterozygous (Qq genotype), while the rest were of undetermined genotype (Figure 2). F2 homozygous (QQ on SIRI0001067) individuals, which were in the top 10% of the population, were used to determine the shared haplotype. Visual examination of all the previously mentioned populations revealed a ∼637-Kb shared haplotype, SIRI0001067, which contained the most significant SNPs (Figure 3). The sharing region contained a total of 16 genes in GenBank. Finkel-Biskis-Jinkins murine osteosarcoma viral oncogene homologue (*FOS*), one of 16 genes, plays essential roles in the osteoclastic differentiation of precursor cells and the upregulation of Receptor Activator of Nuclear Factor κ B (RANK) expression in osteoclast precursors within the bone environment (Arai *et al.* 2012). This gene has been reported to be a candidate for number of vertebrae (Ren *et al.* 2012).

**Figure 2.**
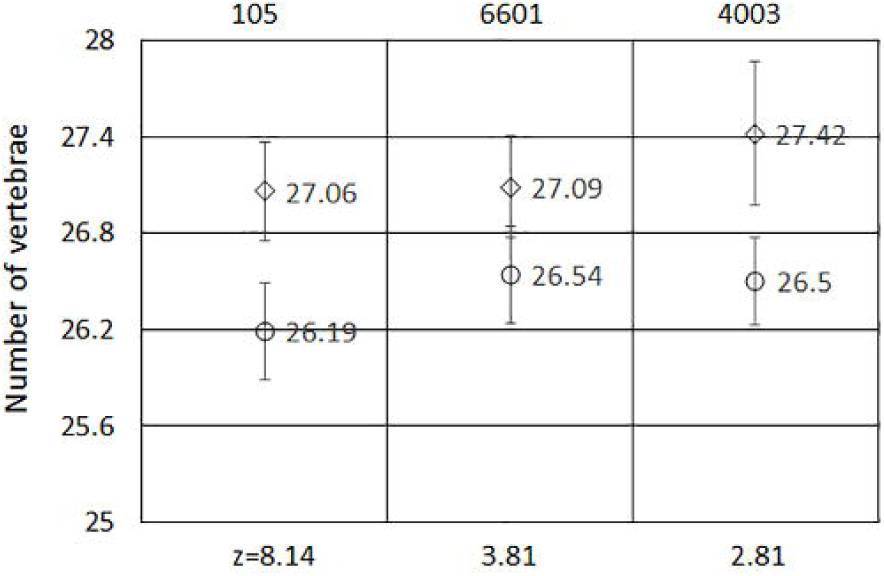
The marker-assisted segregation analysis for F1 boars. The marker-assisted segregation analysis for F1 boars. The graphs show, for 3 F1 boars’ half-sib pedigrees (105, 4003, 6601), the phenotypic mean ± standard errors of the offspring sorted in two groups according to the homolog inherited from the sire. The number of offspring in each group is given above the error bars, respectively. The graph corresponds to the boars that were shown to be heterozygous Qq and reports a Z-score for each pedigree. Q alleles associated with a positive allele substitution effect on number of vertebrae are marked by a diamond, q alleles by a circle. The number within the symbols differentiates the Q and q alleles according to the associated marker genotype.

**Figure 3.**
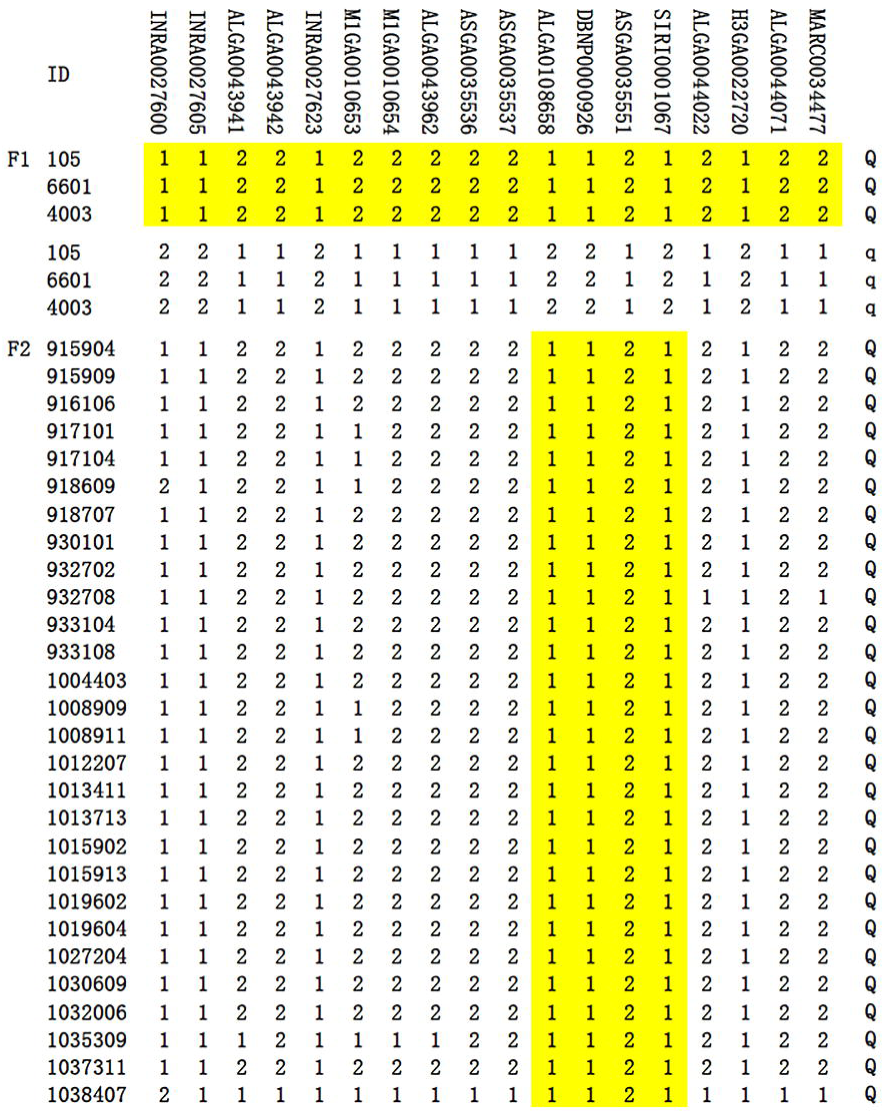
Haplotype sharing analysis in the 4.56-Mb region on SSC7. Shared haplotypes of Qq F1 boars (Large White × Minzhu intercross population) and top 10% F2 individuals with Q chromosomes were analyzed. Polymorphisms are displayed at the respective SNP markers. SNP alleles are shown by 1 and 2 for the major and minor alleles, respectively. Identities of animals carrying the Q chromosome are given in the left axis.

However, the most interesting gene in this region is transforming growth factor, beta 3 (*TGFB3*). This GWAS located the most significant SNP, SIRI0001067, in the intron of *TGFB3*. The development of the vertebral column is a consequence of a segment-specific balance between proliferation, apoptosis, and differentiation of mesoderm cells in embryos (Christ *et al.* 2000). Gene expression analysis shows that *TGFB3* is abundant in the growing undifferentiated mesoderm (Lorda-Diez *et al.* 2010). In zebrafish, *TGFB3* is moderately expressed in 14- to 24-somite embryos (Cheah *et al.* 2005). In mammals, *TGFB3* promotes chondrogenesis in posterfrontal suture-derived mesenchymal cells, influencing different stages of chondrogenic differentiation and proliferation (James *et al.* 2009). *TGFB3* plays a critical role in vertebral column development by increasing the proliferation of mesoderm cells. *TGFB3* protein, in combination with its downstream factor, TGF beta receptor type I (*ALK5*), regulates the differentiation and proliferation of the spinal column (Zhao *et al.* 2014) and has been associated with the biological functions of TGFB3 proteins. Compared with the wild type, Alk5^+/+^ and Alk5^+/-^ mice had an additional 3 and 5 vertebrae, respectively (Andersson *et al.* 2006). Although *TGFB3* knockout mice (TGFB3^-/-^) have been created, they died soon after birth and the effect of gene knockout on number of vertebrae was not recorded (Kaartinen *et al.* 1995). We believe that these GWAS findings support the idea that *TGFB3* might be a strong candidate of the QTL for the number of vertebrae.

In summary, this work is a GWAS focusing on the number of vertebrae in pigs. A genome-wide scan identified 21 significant SNPs within an 8.82-Mb region containing the reported causal gene *NR6A1* on SSC1. An additional 18 SNPs were identified within a 4.56-Mb region on SSC7 that showed genome-wide association with the number of vertebrae. Finally, haplotype sharing analysis refined the 4.56-Mb region to a region about 637 Kb in size, encompassing the gene *TGFB3* on SSC1. Exploration of the gene via additional genetic and functional studies in mammals revealed that *TGFB3* could be a strong candidate for the number of vertebrae in pigs.

## Conflict of Interests

The authors have declared that no conflict of interest exists.

## ACKNOWLEDGMENTS

This research was supported by the Agricultural Science and Technology Innovation Program (ASTIP-IAS02), National Key Technology R&D Program of China (No.2011BAD28B01), earmarked fund for Modern Agro-industry Technology Research System, and Chinese Academy of Agricultural Sciences Foundation (No.2014ZL006).

## LITERATURE CITED

Amin, N., C. M. van Duijn, and Y. S. Aulchenko, 2007 A genomic background based method for association analysis in related individuals. PLoS One 2: e1274.

Andersson, O., E. Reissmann, and C. F. Ibáñez, 2006 Growth differentiation factor 11 signals through the transforming growth factor-beta receptor ALK5 to regionalize the anterior-posterior axis. EMBO Rep. 7: 831–837.

Arai, A., T. Mizoguchi, S. Harada, Y. Kobayashi, Y. Nakamichi et al., 2012 Fos plays an essential role in the upregulation of RANK expression in osteoclast precursors within the bone microenvironment. J. Cell Sci. 125: 2910–2917.

Aulchenko, Y. S., D. J. de Koning, and C. Haley, 2007 Genome-wide rapid association using mixed model and regression: a fast and simple method for genome-wide pedigree-based quantitative trait loci association analysis. Genetics 177: 577–585.

Burgosa, C., P. Latorrea, J. Altarribab, J. A. Carrodeguasc, L. Varona et al., 2014 Allelic frequencies of NR6A1 and VRTN, two genes that affect vertebrae number in diverse pig breeds: A study of the effects of the VRTN insertion on phenotypic traits of a DurocxLandrace-Large White cross. Meat Sci. 100C: 150–155.

Cheah, F. S., E. W. Jabs, and S. S. Chong, 2005 Genomic, cDNA, and embryonic xxpression analysis of zebrafish transforming growth factor beta 3 (tgfbeta3). Dev. Dyn. 232: 1021–1030.

Christ, B., R. Huang, and J. Wilting, 2000 The development of the avian vertebral column. Anat. Embryol. (Berl) 202: 179–194.

Edwards, D. B., C. W. Ernst, N. E. Raney, M.E. Doumit, M.D. Hoge et al., 2008 Quantitative trait locus mapping in an F2 Duroc6Pietrain resource population: II. carcass and meat quality trait. J. Anim. Sci. 86: 254–266.

Fan, Y., Y. Xing, Z. Zhang, H. Ai, Z. Ouyang et al. 2013 A further look at porcine Chromosome 7 reveals *VRTN* variants associated with vertebral number in Chinese and Western Pigs. PLoS One 8: e62534.

Fontanesi, L., A. Ribani, E. Scotti, V. J. Utzeri, N. Veličković et al. 2014 Differentiation of meat from European wild boars and domestic pigs using polymorphisms in the *MC1R* and *NR6A1* genes. Meat Sci. 98: 781–784.

Hirose, K., S. Mikawa, N. Okumura, G. Noguchi, K. Fukawa et al. 2013 Association of swine *vertnin* (VRTN) gene with production traits in Duroc pigs improved using a closed nucleus breeding system. Anim. Sci. J. 84: 213–221.

James, A. W., Y. Xu, J. K. Lee, R. Wang, and M. T. Longaker, 2009 Differential effects of TGF-beta1 and TGF-beta3 on chondrogenesis in posterofrontal cranial suture-derived mesenchymal cells in vitro. Plast. Reconstr. Surg. 123: 31–43.

Kaartinen, V., J. W. Voncken, C. Shuler, D. Warburton, D. Bu et al. 1995 Abnormal lung development and cleft palate in mice lacking TGF-beta 3 indicates defects of epithelialmesenchymal interaction. Nat. Genet. 11: 415–421.

King, J. W. B., and R. C. Roberts, 1960 Carcass length in the bacon pig: its association with vertebrae numbers and prediction from radiographs of the young pig. Anim. Prod. 2: 59–65.

Lorda-Diez, C. I., J. A. Montero, J. A. Garcia-Porrero, and J. M. Hurle, 2010 Tgfbeta2 and 3 are coexpressed with their extracellular regulator Ltbp1 in the early limb bud and modulate mesodermal outgrowth and BMP signaling in chicken embryos. BMC Dev. Biol. 10: 69.

Madsen, P., P. Sørensen, G. Su, L. H. Damgaard, H. Thomsen et al. 2006 DMU-a package for analyzing multivariate mixed models. In 8th World Congress on Genetics Applied to Livestock Production, Brasil.

Mikawa, S., T. Hayashi, M. Nii, S. Shimanuki, T. Morozumi et al. 2005 Two quantitative trait loci on Sus scrofa chromosones 1 and 7 affecting the number of vertebrae. J. Anim. Sci. 83: 2247–2254.

Mikawa, S., T. Morozumi, S. Shimanuki, T. Hayashi, H. Uenishi et al. 2007 Fine mapping of a swine quantitative trait locus for number of vertebrae and analysis of an orphan nuclear receptor, germ cell nuclear factor (NR6A1). Genome Res. 17: 586–593.

Mikawa, S., S. Sato, M. Nii, T. Morozumi, G. Yoshioka et al. 2011 Identification of a second gene associated with variation in vertebral number in domestic pigs. BMC Genet. 12: 5.

Miller, S. A., D. D. Dykes, and H. F. Polesky, 1988 A simple salting out procedure for extracting DNA from human nucleated cells. Nucleic Acids Res. 16: 1215.

Nezer, C., C. Collette, L. Moreau, B. Brouwers, J. J. Kim et al. 2003 Haplotype sharing refines the location of an imprinted quantitative trait locus with major effect on muscle mass to a 250-kb chromosome segment containing the porcine IGF2 gene. Genetics 165: 277–285.

Ren, D. R., J. Ren, G. F. Ruan, Y. M. Guo, L. H. Wu et al. 2012 Mapping and fine mapping of quantitative trait loci for the number of vertebrae in a White Duroc × Chinese Erhualian intercross resource population. Anim. Genet. 43: 545–551.

Rubin, C. J., H. J. Megens, A. Martinez Barrio, K. Maqbool, S. Sayyab et al. 2012 Strong signatures of selection in the domestic pig genome. Proc. Natl. Acad. Sci. U. S. A. 109: 19529–19536.

Sato, S., Y. Oyamada, K. Atsuji, K. Atsuji, T. Nade, S. Sato et al. 2003 Quantitative trait loci analysis for growth and carcass traits in a Meishan × Duroc F2 resource population. J. Anim. Sci. 81: 2938–2949.

Wada, Y., T. Akita, T. Awata, T. Furukawa, N. Sugai et al. 2000 Quantitative trait loci (QTL) analysis in a Meishan × Göttingen cross population. Anim. Genet. 31: 376–384.

Yang, G., J. Ren, Z. Zhang, and L. Huang, 2009 Genetic evidence for the introgression of Western NR6A1 haplotype into Chinese Licha breed associated with increased vertebral number. Anim. Genet. 40: 247–250.

Yang, Q., J. Cui, I. Chazaro, L. A. Cupples, and S. Demissie, 2005 Power and type I error rate of false discovery rate approaches in genome-wide association studies. BMC Genet. 6: S134.

Zhang, L. C., N Li., X. Liu, J. Liang, H. Yan et al. 2014 A genome-wide association study of limb bone length using a Large White□×□Minzhu intercross population. Genet. Sel. Evol. 46: 56.

Zhang, Z. G., B. D. Li, and X. H. Chen, 1986 Pig breeds in China. Shanghai Scientific and Technical Publisher, Shang, China.

Zhao, S. F., M. Z. Chai, M. Wu, Y. H. He, T. Meng et al. 2014 Effect of vitamin B12 on cleft palate induced by 2,3,7,8-tetrachlorodibenzo-p-dioxin and dexamethasone in mice. J. Zhejiang Univ. Sci. B 15: 289–294.

